# An inducible CRISPR-interference library for genetic interrogation of *Saccharomyces cerevisiae* biology

**DOI:** 10.1101/2020.03.05.978619

**Authors:** Amir Momen-Roknabadi, Panos Oikonomou, Maxwell Zegans, Saeed Tavazoie

## Abstract

Genome-scale CRISPR interference (CRISPRi) is widely utilized to study cellular processes in a variety of organisms. Despite its dominance as a model eukaryote, a genome-wide CRISPRi library, optimized for targeting the *Saccharomyces cerevisiae* genome, has not been presented to date. We have generated a comprehensive, inducible CRISPRi library, based on spacer design rules optimized for yeast. We have validated this library for genome-wide interrogation of gene function across a variety of applications, including accurate discovery of haploinsufficient genes and identification of enzymatic and regulatory genes involved in adenine and arginine biosynthesis. The comprehensive nature of the library also revealed parameters for optimal transcriptional repression, including upstream distance, nucleosomal occupancy, and strand bias. CRISPRi screens, using this library can identify genes and pathways with high precision and low false discovery rate across a variety of experimental conditions, enabling rapid and reliable genome-wide assessment of gene function and genetic interactions in *S.cerevisiae*.

## Introduction

Technologies that generate systematic genetic perturbations have revolutionized our ability to rapidly determine the genetic basis of diverse cellular phenotypes and behaviors (1–7). CRISPR-Cas9 technology (8–10), due to its high-fidelity and relatively low off-target effect, has become the dominant method for systematic, high-throughput genetic screening in diverse eukaryotic systems (11–14). Nuclease-deactivated Cas9 (dCas9) has facilitated genome-wide screens further by enabling transient modulation of target gene expression (15,16). This is achieved by fusing dCas9 to an inhibitory or activating domain to repress (CRISPRi) or activate (CRISPRa) gene expression (17–22). At the same time, the use of array-based oligonucleotide synthesis has enabled production of large-scale spacer libraries for use in genome-wide applications.

One complicating factor in the use of CRISPRi technology is balancing the efficacy of targeting with limiting the off-target activity of CRISPR/dCas9 machinery. Hence, many studies have aimed to determine the rules for efficient gRNA design (16–19). For example, in K562 human myeloid leukemia cells, optimal gRNAs are found to target a window of −50 bp to +300 bp relative to the transcription start site (TSS) of a gene (23). In *Saccharomyces cerevisiae*, however, the ideal guide positioning differs from human cell lines. Smith *et al.* (22) found that the optimal window is a 200 bp region immediately upstream of the TSS. The same group in a subsequent study refined their earlier findings showing that the region between TSS and 125 bp upstream of TSS is more effective for CRISPR-mediated repression (24). In addition, they showed a positive correlation between guide efficiency and chromatin accessibility scores, bolstering the notion that location of target in relation to TSS is not the only determinant of gRNA efficacy (22). The positioning and design rules of gRNAs are therefore organism specific. Recently, Lian et al (25) developed a multi-functional genome-wide CRISPR (MAGIC) system for high throughput screening in *S.cerevisiae*. Although they successfully used a combinatorial approach to map the furfural resistance genes, their system did not utilize yeast-specific design rules (22). More importantly, their CRISPR system is not inducible making it difficult to perform context-dependent repression of gene function, and to survey the role of essential genes in arbitrary phenotypes of interest.

Here, we introduce an inducible genome-scale library, dedicated for CRISPRi in *S. cerevisiae*, and designed based on the rules described by Smith *et al* (22,24). We demonstrate the efficacy of this library in targeting essential genes and identifying dosage-sensitive ORFs. In addition, the library enabled us to identify genes involved in Adenine and Arginine biosynthesis using only a single round of selection. This CRISPRi library and protocol can, thus, be used to efficiently and inexpensively perform genome-wide knock-down screens to discover the genetic basis of any selectable phenotype. In addition, the ability to easily perform CRISPRi screens in a desired genetic background of interest, enables rapid profiling of genetic interactions between a desired allele and knock-downs of all the genes in the genome.

## Results and Discussion

### Design and Construction of a Whole-Genome CRISPRi Library

We developed our CRISPRi library largely based on the design principles of Smith *et al*. (22). Smith *et al*. (22) designed a single-plasmid inducible system expressing the catalytically inactive *Streptococcus pyogenes* Cas9 (dCas9) fused to the MXI1 transcriptional repressor (17), as well as a gRNA. The gRNA is under the control of a tetO-modified RPR1 RNA polymerase III promoter regulated by a tetracycline repressor (tetR), which is also expressed by this plasmid (18,26). Therefore, the expression of the gRNA is induced by addition of anhydrotetracycline (ATc) to the growth medium. In addition, a NotI site between the tetO and the gRNA scaffold enables the rapid cloning of spacers. TetR and dCas9-Mxi1 are expressed from the GPM1 and TEF1 promoters, respectively. For compatibility with an ongoing project in our group, we have replaced the *URA3* selection marker in PRS416 with *HIS3*. We call this plasmid amPL43 (Figure 1a). In order to validate the effectiveness of our system, we cloned gRNAs targeting *ERG25*, *ERG11* and *SEC14* genes in amPL43. Upon addition of ATc, these genes were repressed when compared to the samples without ATc (per qRT-PCR, Figure 1b). The repression was seen as early as one hour after induction and could reach as much as 10-fold over 24 hours, depending on the target.

**Figure 1.**
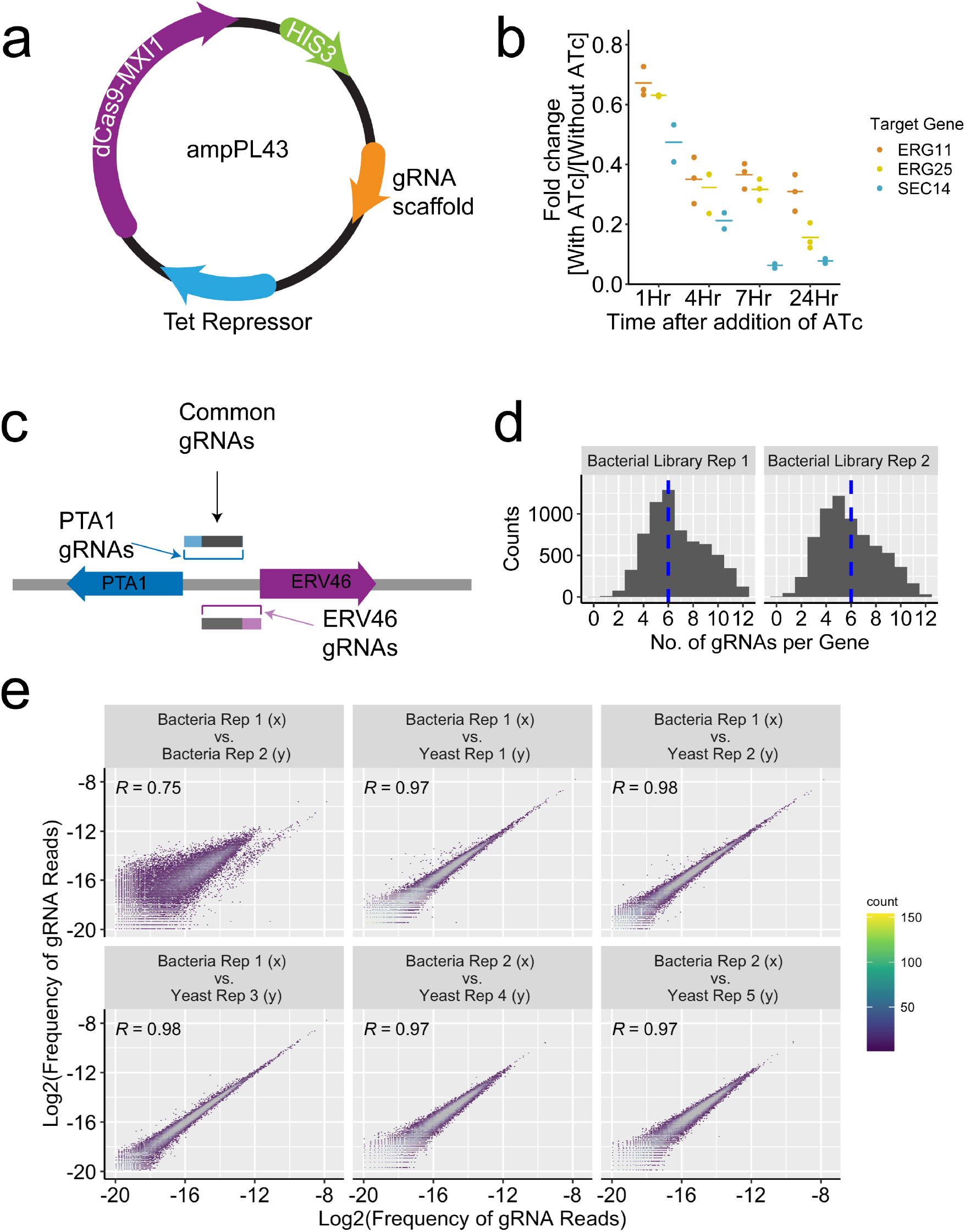
**a)** Schematic of amPL43 expression vector for inducible CRISPRi library in *S. cerevisiae*. gRNA scaffold region contains a NotI site for cloning gRNA sequences, an RPR1 promoter with a TetO site, and a constant region for the gRNA. **b)** The expression fold-change of each target: ERG25 (three replicates), ERG11 (three replicates), and Sec14 (two replicates), as a result of gRNA induction by ATc, was calculated over time by qPCR. The mean for each sample is represented by a solid line. **c)** Schematic depicting the genomic region of PTA1 and ERV46. The gRNAs targeting the region between the two genes, depending on their proximity to each gene, could affect both genes. **d)** Histogram depicting the number of gRNAs per gene in two library replicates. The dashed blue line denotes the median: ~6. **e)** Scatter plot depicting the frequency of reads per gRNA between select biological replicates of the CRISPRi library. Pearson Correlation R value is reported for each pair.

After confirming the effectiveness of this mode of CRISPR-inhibition, we constructed a genome-wide CRISPRi library to target all *S. cerevisiae* genes. To this end, we obtained and ranked all possible spacer sequences targeting every open reading frame (ORF) based on their distance to TSS and nucleosome score (22,24). We then selected the top six gRNAs for each ORF. The dCAS9-MXI1 mediated repression could affect the genes on both plus and minus strands (18). Therefore, some of the selected sequences could be targeting a neighboring gene with a shared intergenic region. For example, a portion of the gRNAs targeting PTA1 could affect ERV46 (Figure 1c). For genes that share a gRNA with another gene, we selected up to six additional sequences unique to those genes (See Methods). In order to evaluate the guide design parameters, we included ~5,000 gRNAs that target further upstream of TSS or downstream of TSS. Altogether, we designed more than 51,000 gRNAs, with between 6-12 sequences per gene (Supplementary table 1). To generate a negative control set of gRNAs, we also included 500 synthetic randomly shuffled gRNAs with no matches in the yeast genome (Supplementary table 2).

This oligonucleotide library was synthesized on a 92918-format chip (CustomArray, Bothell, WA, USA) and cloned into amPL43 using universal adapter sequences. Briefly, the pooled oligonucleotides were amplified by PCR, cloned using Gibson Assembly (27), and transformed into DH5α *E.coli* (NEB C2987H, >100x colonies/gRNA). The transformed bacteria were grown in a semisolid LB as individual colonies (0.35% Seaprep Agarose, Lonza, Basel, Switzerland) to minimize competition between the strains. Semisolid 3D media provides a more cost-effective and less labor-intensive method for large-scale libraries than conventional 2D plating. The pooled plasmid library was transformed into BY4741 using the standard LiAc/PEG method (28) with minor modifications, and grown in semisolid SC-His+glu media (0.35% Seaprep Agarose, two or three biological replicates) for 48 hours, pooled, and resuspended in SC-His+glu media, and frozen for future use.

The short gRNA sequences can act as unique identifiers of individual strains and, like barcodes, can be quantified using next-generation amplicon sequencing. To this end, the plasmids were extracted from yeast samples, and the targeting region was PCR amplified with flanking Illumina adapters and submitted for sequencing. We generated two independently cloned bacterial library replicates. At the realized sequencing depth, our first library replicate included more than 41,000 gRNAs, targeting all of the yeast ORFs except YBR291C, while more than 36,000 gRNAs were present in the second CRISPRi library replicate, targeting all but five of the yeast ORFs (YBL023C, YIL171W, YJL219W, YOR324C, YPR019W, Figure 1d and Supplementary Figure 1). The two bacterial libraries are highly correlated (Figure 1e, Pearson correlation R=0.75). The loss in the number of gRNAs after cloning could be attributed to limited sequencing depth (~7 million reads for the first replicate and ~4 million reads for the second replicate) or synthesis errors. Biological replicates from the two bacterial libraries were transformed into yeast. There is a strong correlation between the frequency of reads of the gRNAs in the bacterial plasmid libraries and those transferred to yeast demonstrating that there is no systemic bias as a result of the transformation. (Figures 1e, Supplementary Figure 2). The five separate biological replicates of yeast library gave highly reproducible diversity and abundance (Supplementary Figure 3 and Supplementary Table 3). The gRNA library does not show a bias for any specific GO Term and shows good representation across compartments, functions and biological processes (Supplementary Figure 4).

### High-Throughput Identification of Dosage sensitive Genes

To demonstrate the utility of our CRISPRi library for high-throughput genotype-phenotype mapping, we set out to determine whether we could systematically discover dosage-sensitive genes by using a simple outgrowth experiment. We focused on determining how well these dosage-sensitive genes corresponded to genes previously found to be haploinsufficient in yeast (2). One of the major advantages of our library is its inducibility. Inducible gRNA expression allows us to efficiently target dosage-sensitive genes for short intervals and determine their phenotypic outcome. To this end, we inoculated semisolid media (to avoid direct competition between strains) with distinct gRNA targets. We introduced the equivalent of 1 OD_660_ of the library (with an average of ~450 copies of each library member) in media with and without ATc induction. The fitness consequence of knocking down every yeast gene can be determined using the depletion or enrichment of the barcodes in the library in a comparison of induced and un-induced samples. Under these conditions, we expect that the gRNAs targeting haploinsufficient genes should be significantly depleted in the induced library (ATc+).

We grew three library replicates for 24 hours, extracted plasmids from pooled samples, and prepared amplicons for NGS (See Methods). We calculated the log-fold change of reads between samples with and without ATc, for these replicates. To calculate the statistical significance of gRNA depletion, we simulated a library of synthetic scrambled genes by sampling the scores from the synthetic randomly shuffled gRNAs to create a baseline (see methods for details). Not all gRNAs will efficiently repress the expression of their target gene (22). Therefore, we only focused on the impact of the most effective gRNAs for a given gene. To this end, we sorted the gRNAs for each gene based on their ratio and designated the mean of the most effective three gRNAs as the score for that gene (See Methods). We used the synthetic scrambled gene distribution to calculate Z-scores from the gene depletion scores for each replicate. Next, we averaged the Z-scores for the biological replicates using Stouffer's method. The gene depletion scores between the replicates were well correlated (Figure 2a). This correlation is much stronger for the haploinsufficient genes, demonstrating that the repression as the result of the gRNA induction is reproducible (Figure 2b). The gRNAs targeting known haploinsufficient genes are significantly depleted in the induced compared to the uninduced sample (*p-*value<1×10^−15^ Wilcoxon signed-rank test). We used the receiver operating characteristic (ROC) curve analysis to find the threshold, measure FDR, and determine which genes are significantly affected as the result of gRNA induction at a given FDR. With only a single round of growth selection, and three replicates, we were able to correctly categorize ~85% of haploinsufficient genes with FDR<10% or 81% with FDR of <5% (Figures 2c-d and Supplementary Tables 4-5).

**Figure 2.**
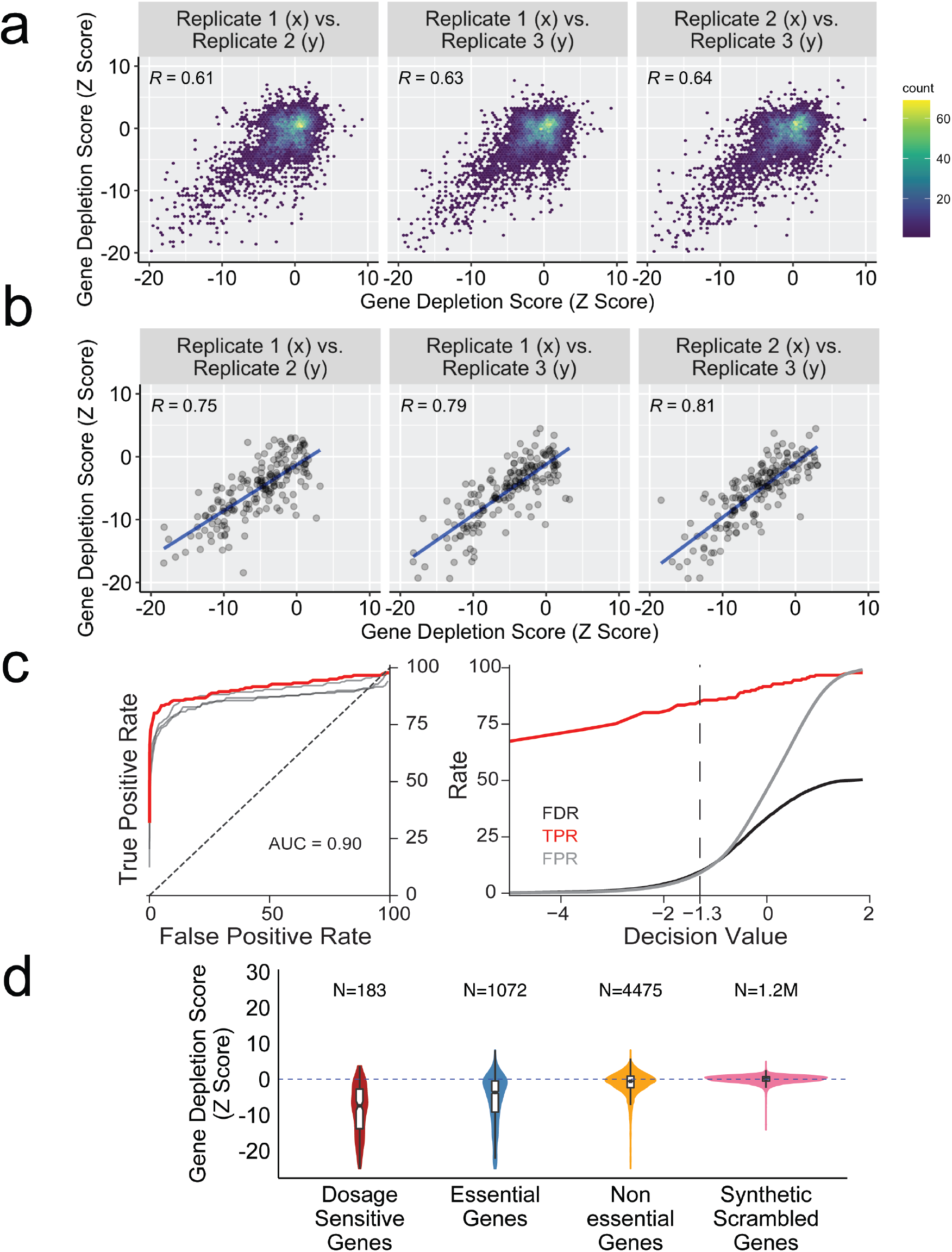
**a)** Binned scatter plots of gene depletion score between the replicates. **b)** Binned scatter plots of gene depletion score, limited to haploinsufficient genes, between the replicates. **c)** The Receiver Operating Characteristic Curve (ROC) for the detection of haploinsufficient genes and FDR, TPR and FPR trends based on decision values. ROC curve shows that the depletion Z Score is a strong classifier for haploinsufficient genes. The individual replicates are shown in grey. Area under the curve is 0.90. Dashed line denotes the decision value for FDR<10%. **d)** Violin plots depicting the gene depletion score distribution for the haploinsufficient genes, essential genes, nonessential genes and synthetic scrambled genes, resulting from 200x random sampling of synthetic randomly shuffled gRNAs, in induced (SC-His +ATc) versus uninduced samples (SC-His-ATc).

A major advantage of our inducible CRISPRi system is the ability to interrogate the role of essential genes in any selectable phenotype. In order to explore this capacity, we set out to determine what percentage of known essential genes are also dosage sensitive under our experimental conditions. The sensitivity and specificity of CRISPRi-based discovery of essential genes are limited by fundamental biological factors. On one hand, although essential genes are associated with functions that are indispensable to cellular life, it has been shown that not all essential genes are dosage sensitive (2). This implies that even reducing the dosage of some essential genes to 50% would not measurably affect cellular fitness. On the other hand, essential genes are overrepresented among dosage-sensitive genes (2). Therefore, it is of interest to determine what percentage of known essential genes show dosage-sensitivity and therefore can be detected by systematic CRISPRi knock-down. Indeed, we observed that the majority of essential genes exhibit significantly lower gene depletion scores (*p-* value<1×10^−15^, Wilcoxon signed-rank test), while the non-essential genes were largely unaffected (Figure 2d). Overall, as is shown in the ROC curve analysis (Supplementary Figure 5a), we observed that ~67% of essential genes (FDR<10%) show dosage-sensitivity based on their gene depletion scores.

The genome wide nature of our library, and its sensitivity to identify dosage-sensitive genes, enabled us to determine whether there was spatial bias for effective targeting by gRNAs. In particular, we were interested in factors such as strand, distance to TSS, and nucleosome occupancy score. As can be seen in Figure 3a, gRNAs located between TSS and 150 bp upstream of TSS are particularly effective for detecting dosage sensitive essential genes, with the strongest effect for the guides targeting 50 bp upstream of TSS, thus further refining yeast CRISPRi design rules (24). In addition, as may be expected, a higher nucleosome score seems to reduce the effectiveness of gRNA mediated transcriptional repression, with the most effective gRNAs having a nucleosome score near zero (Figure 3b). As expected, gRNAs targeting nonessential genes did not show any dependency between the gRNA depletion score and the distance to TSS or the nucleosome occupancy score (Figure 3c-d). Finally, we investigated whether gRNAs’ effectiveness is influenced by the strandedness of gRNA targeting. To minimize the effect of other factors, we only focused on gRNAs targeting dosage-sensitive essential genes that have a nucleosome occupancy score less than 0.1 and are within 125 bp upstream of TSS. The mean of the gRNA depletion score for gRNAs with the PAM on the same strand as the ORF was −1.32 while the mean for the gRNAs on the opposite strand was −0.93 (Figure 3e, Wilcoxon signed-rank test *p-*value <8.8×10^−8^). This indicates that, while gRNAs can target and repress ORFs on both strands, the strandedness significantly influences gRNAs’ effectiveness. In addition, the gRNAs with the PAM on the same strand as TSS are most effective when located between 50 to 75 bp upstream of the TSS, while the gRNAs with the PAM on the opposite strand, have a maximum efficacy between 25 to 50 bp upstream of the TSS (Figure 3f-g).

**Figure 3.**
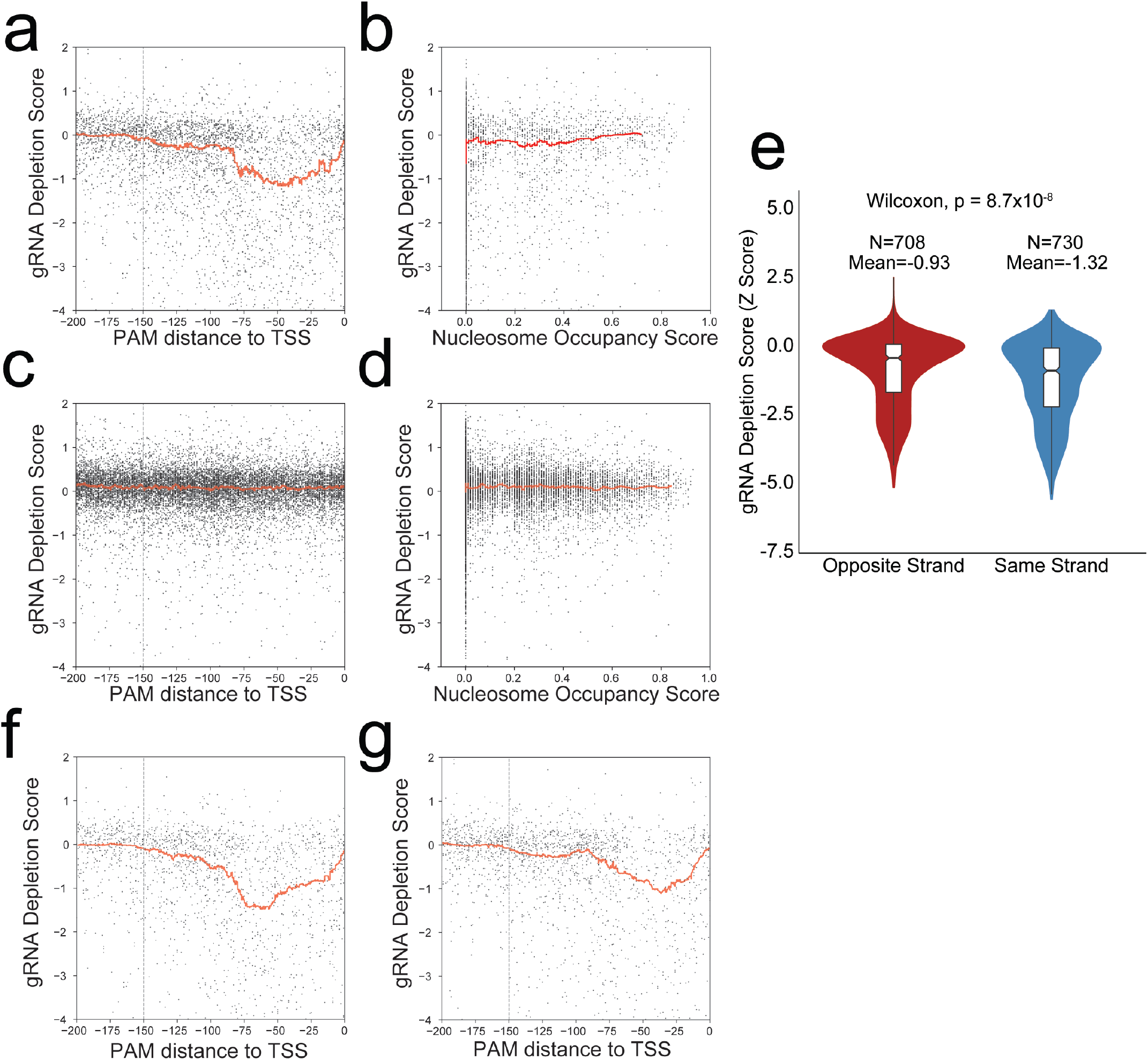
**a)** Scatter plot depicting the gRNA depletion score for dosage sensitive essential genes versus the distance from TSS. The red line represents the rolling average (window of 200). The dashed line signifies 150 bp upstream of TSS. **b)** Scatter plot depicting the gRNA depletion score for dosage sensitive essential genes versus the nucleosome occupancy score. The red line represents the rolling average (window of 200). **c)** Scatter plot depicting the gRNA depletion score for nonessential genes versus the distance from TSS. The red line represents the rolling average as above. The dashed line signifies 150 bp upstream of TSS. The red line represents the rolling average as above. **d)** Scatter plot depicting the gRNA depletion score for nonessential genes versus the nucleosome occupancy score. The red line represents the rolling average as above. **e**) Violin plots depicting the distribution of gRNA depletion scores for gRNAs targeting the opposite or same strand as the target ORF. **f)** Scatter plot depicting the gRNA depletion score for gRNAs targeting dosage sensitive essential genes with PAM on the same strand as TSS versus the distance from TSS. The red line represents the rolling average as above. The dashed line marks 150 bp upstream of TSS. **g)** Scatter plot depicting the gRNA depletion score for gRNAs targeting dosage sensitive essential genes with PAM on the opposite strand as TSS versus the distance from TSS. The red line represents the rolling average as above. The dashed line marks 150 bp upstream of TSS.

### Identification of the Genes Involved in Biosynthesis of Adenine and Arginine

Next, we explored whether our CRISPRi based approach can efficiently identify smaller sets of genes associated with specific biological processes. We, thus, chose to identify the genes involved in two distinct biosynthetic pathways, Arginine and Adenine. This was done by CRISPRi profiling, across five replicates, of induced cells grown in drop-out media for arginine and adenine, compared to cells grown in media including the nutrients. Histidine is absent in both conditions as a necessary condition for plasmid maintenance. We expected gRNAs targeting genes that contribute to general cellular fitness to be depleted to a similar degree for both samples, while gRNAs involved in the biosynthesis of arginine and adenine would be differentially affected. As such, we quantified the depletion of gRNAs in the arginine/adenine/histidine drop out media against the histidine drop out media control. In order to systematically determine all the pathways that were affected, we used iPAGE (29), a sensitive pathway analysis tool that directly quantifies the mutual information between pathway membership and the global distribution of gene depletion scores. The iPAGE results show that the biological processes for Arginine biosynthesis and purine biosynthetic processes (30–36) are significantly informative of the depletion of gRNAs targeting the pathways (Supplementary Figure 6 and Supplementary Tables 4-5). In addition, the gRNAs targeting the general pathways for ATP export, nitrogen starvation and amino acid biosynthesis were also depleted. The iPAGE analysis also showed that gRNAs targeting pathways affecting protein dynamics, such as translation and import of proteins into the nucleus, are significantly enriched. This suggests that the activity of these pathways partially affect growth fitness when cells are simultaneously deprived of arginine and adenine. We also detected significant depletion/enrichment of pathways associated with protein sorting, such as endosome to Golgi transport and retrograde vesicle transport. This is in line with previous observations that mutations in many of the arginine biosynthesis genes are known to cause abnormal vacuole morphology (37). Our data, thus, provides additional evidence that arginine deficiency undermines protein sorting functions in *S. cerevisiae*.

Our screen detected 11 out of 19 genes annotated in core arginine biosynthesis and adenine biosynthesis pathways (Figure 4a-b, Supplementary Figure 5b, FDR<10%). Arginine biosynthesis is a particularly complex biosynthetic pathway with connections to several other pathways, such as polyamine and pyrimidine biosynthesis, and certain degradative pathways (32,38). In addition, transcription of arginine biosynthetic genes is repressed in the presence of arginine by the ArgR/Mcm1p complex, which consists of Arg80p, Arg81p, Arg82p, and Mcm1p (39). As such, downregulation of the ArgR/Mcm1p complex genes would be expected to increase the fitness of the cell in the absence of arginine in the media. Consistent with this expectation, we found that the gene depletion scores for these genes were positive, providing further support for the sensitivity of our CRISPRi screen to detect positive and negative contributors to a phenotype of interest (Figure 4a). Our analysis of gRNA efficiency showed that the gRNAs with PAM 0 to 150 bp upstream of each TSS are particularly effective. Therefore, we explored whether we could improve the precision of our detection by limiting our analysis to gRNAs that target 0-150 bp upstream of the TSS, while maintaining gRNA diversity. As is shown in Figure 4c, we detected 15 out of 19 adenine and arginine biosynthesis genes (FDR<10%, Supplementary Table 4).

**Figure 4.**
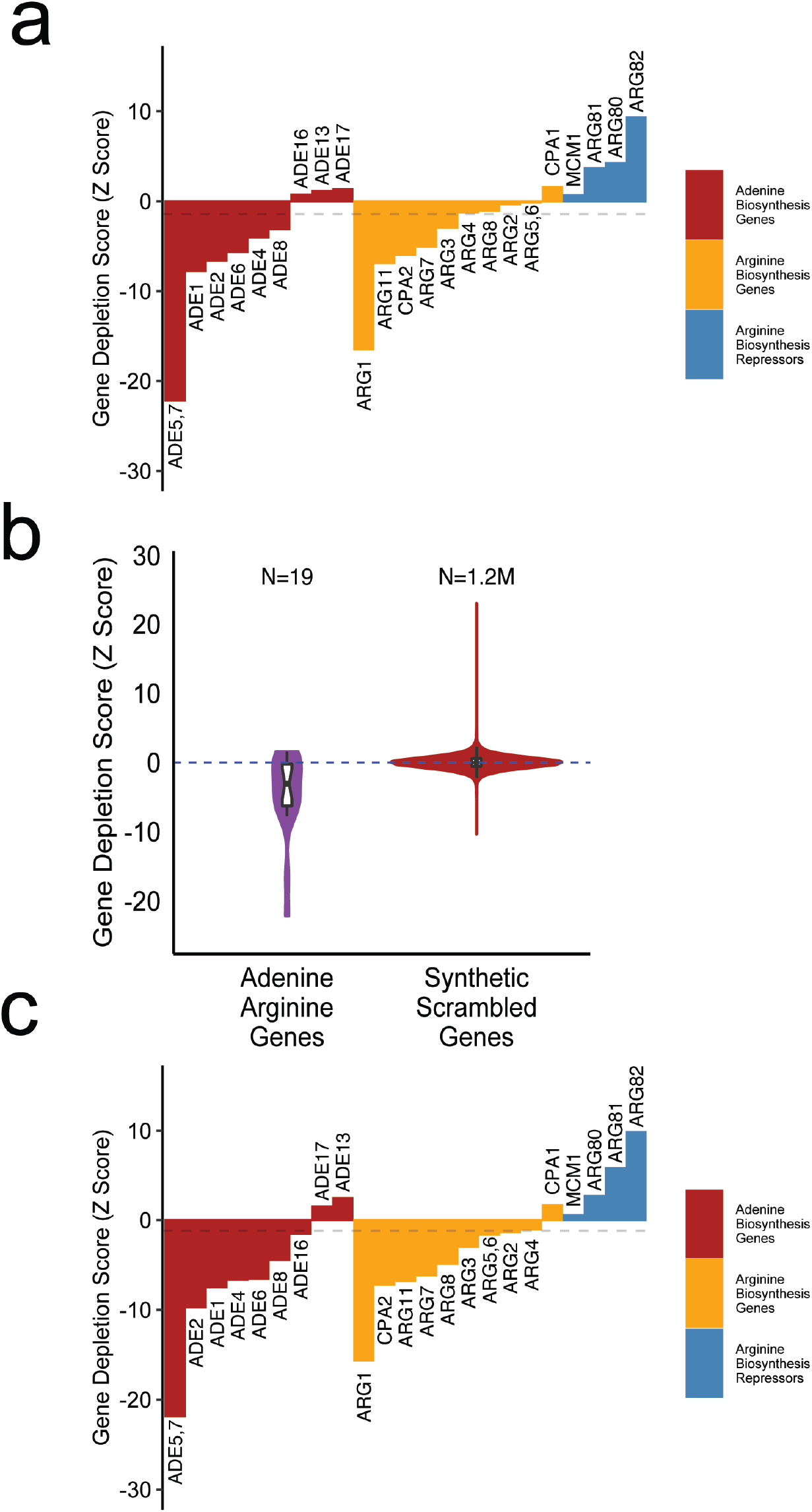
**a)** Bar plots showing the depletion scores for the known annotated genes involved in adenine and arginine biosynthesis pathways in addition to arginine regulatory genes. Dashed gray line marks the threshold for FDR= 10%. **b)** Violin plots depicting the gene depletion score distribution for the adenine arginine deprivation experiment (SC-His-Ade-Arg +ATc vs. SC-His +ATc), shown for the known Adenine/Arginine Biosynthetic genes, and synthetic scrambled genes, resulting from 200x random sampling of synthetic randomly shuffled gRNAs. **c)** Bar plots showing the depletion scores for the annotated genes involved in adenine and arginine biosynthesis pathways in addition to arginine regulatory genes, restricted to gRNAs with PAM located within 150 bp of TSS. Dashed gray line marks the threshold for FDR= 10%.

It is important to point out that our FDR estimate of 10% is conservative, since strict gene ontology membership of these pathways does not capture the full complexity of the highly interactive genetic and regulatory networks coordinating nucleotide and amino-acid metabolism. For example, in addition to the genes involved in arginine and adenine biosynthesis, most depleted genes were involved in protein sorting and other general amino acid metabolic pathways such as SSY1 which is known to sense external amino acid levels (40).

## Discussion

Here, we have established a versatile genome-wide functional screening library for CRISPRi in *S. cerevisiae* and further refined the gRNA design rules for efficient transcriptional repression. The ability to interrogate essential genes, led us to discover that 67% of essential genes also exhibit dosage sensitivity under our experimental conditions. Our focused study of arginine/adenine deprivation demonstrates attractive sensitivity/specificity characteristics for probing the genetic basis of arbitrary phenotypes and biological processes. This CRISPRi library has distinct advantages compared to current available genome-wide methods: The gRNAs are specifically designed based on *S. cerevisiae* specific rules, and more importantly, the repression is inducible, enabling control over the scale, context, and timing of gene perturbations. The ability to quantitatively probe the role of essential genes is also a major advantage of inducible CRISPRi over both CRISPR and gene-deletion library approaches. In addition, our use of 3D semisolid agarose to generate and interrogate large diverse libraries provides a more efficient approach over traditional 2D plating protocols while, at the same time, minimizing competitive biases that confound liquid outgrowth. Furthermore, the ability to easily transform the library into any genetic background of interest will enable rapid, parallel mapping of genetic interactions for any allele of interest (41). These advantages make our CRISPRi library a powerful and versatile tool for genetic interrogation of yeast biology.

## Material and Methods

### Strains, Plasmids and Media

PRS416-MXI1 was ordered from AddGene (Cambridge, MA, USA). In order to create amPL43, the PRS416-MXI1 backbone was PCR amplified. His3 was amplified from pSH62 (42). The primers are listed in Supplementary Table 6. Gibson assembly (NEB, Ipswich, MA, USA) was used to create amPL43. The plasmid’s sequence was verified using Sanger sequencing. gRNA oligo library was purchased from Custom Array (92918-format chip, Bothell, WA, USA). The library spacers were amplified and extended as described by Smith *et al.* (22). To generate the library, amPL43 was maxiprepped (Qiagen, Hilden, Germany) and cut with NotI. Gibson Assembly was used to clone the oligos in the NotI site of amPL43. The library was transformed into DH5α *E.coli* (NEB C2987H). The transformed bacteria were grown in 1L of semisolid LB plus 100 μg/ ml ampicillin (0.35% Seaprep Agarose, Lonza) to minimize competition between the colonies. Use of semisolid media would minimize the competition and ensure that the growth of the colonies would be independent of each other and avoid any potential jackpotting effects. We sequenced 32 individual colonies to assess the quality of our cloning method. We did not detect any empty vector or repeat sequences. To pool the library, the semisolid media was stirred for 10 minutes using a magnetic stirrer. 50 ml of the library was collected by centrifugation. The pooled library was miniprepped (Promega, Madison, WI, USA) according to manufacturer’s instructions. Two independent library replicates were generated.

Competent BY4741 cells were produced using the standard LiAc/PEG method (28), with minor modifications: 5mL of SC+glucose media was inoculated with a fresh colony of BY4741 and grown overnight in 30°C shaker. 1 ml of over-night culture (OD_660_~ 6) was added to 25 mL of SC+glucose and shaken in 30°C shaker for 4 hours (OD_660_~0.7). Cells were pelleted by centrifugation at 3000g for 5 minutes and washed twice in 25 mL sterile water. Next, cells were resuspended in 1 mL sterile water and transferred to 1.5 mL Eppendorf tube and centrifuged for 1 min at 4000g and then resuspended in sterile water to a final total volume of 1 mL. 200 μL of competent cells were aliquoted into an Eppendorf tube and centrifuged for 30 seconds at 4000g and the supernatant was removed. Next, 240 μL of freshly made sterile 50 % PEG 3500, 36 μL of sterile 1 M lithium acetate, 50 μL of Salmon sperm DNA (Thermofisher, Waltham, MA, USA), and 2 μg of plasmid DNA was added to the cells and the volume was adjusted to 350 μL using sterile water. Then the mixture was vortexed and incubated at 42° C for 20 minutes, vortexed, and again incubated at 42°C for another 20 minutes. Transformed cells were centrifuged and supernatant was removed, and 1 mL of SC+glucose was added to it. The cells were moved to a round bottom falcon tube and shaken at 30°C for 1 hour.

We transformed three yeast replicates from the first bacterial library and two replicates from the second bacterial library. The transformed library was grown in semisolid SC-His+glu media (0.35% Seaprep Agarose) for 48 hours, pooled, and resuspended in SC-His+glu media plus 20% glycerol and frozen for future use. To extract the plasmids, 50 OD_660_ of cells were pooled and resuspended in 2 ml SE buffer (0.9M sorbitol, 0.1M EDTA pH 8.0), 20 μl Zymolyase 100T (2 mg/ml) and 20 μl β-mercaptoethanol and incubated at 37°C for one hour, followed by standard mini prep extraction per manufacturer’s instructions (Promega). The pooled plasmid library was suspended in 50 μL of sterile water. For this study, we inoculated 250ml of the following three semisolid media with a number of cells equivalent to 1 OD_660_ unit of the library: A) SC-His with ATc, B) SC-His-Arg-Ade with ATc C) SC-His-Arg-Ade without ATc. Haploinsufficient gene were derived from Deutschbauer *et al.*, 2005. Nonessential and essential gene lists were derived from Giaever *et al.*, 1999.

### qPCRS

Strains were typically grown in SC-HIS overnight, diluted to an OD_660_ of 0.07 with/without 250 ng/mL ATc. We selected a mix of three different gRNAs per target gene. The sequences for the selected gRNAs are reported in supplementary table 6. The cells were grown at 30°C and a sample was taken from each tube at 1, 4, 7 and 24hrs. RNA was extracted using Norgen Biotek Total RNA Purification Kit (Norgen Biotek, ON, Canada) and cDNA was made using Maxima H First Strand cDNA Synthesis Kit, with dsDNase (Thermofisher). ERG25, ERG11 and Sec14 primers and their corresponding spacers were adapted from Smith *et al.* (22). qRT-PCR was performed using SYBR® Green PCR Master Mix (Life Technologies, Carlsbad, CA, USA) and the Quantstudio 5. ΔΔCt of the target genes’ in induced versus uninduced states as compared to ACT1 level are reported.

### Design of the gRNA Library

The gRNA sequences and their relative location to transcriptional start site (TSS), as well as the nucleosome occupancy scores, were adapted from Smith *et al.* (22). When possible, the gRNAs that were located between 0 and 200 bp of the TSS, were sorted based on their nucleosome occupancy score and the top six gRNAs were chosen. In rare cases, when six gRNAs were not available for any given gene, we searched for the gRNAs further from TSS. When more than six acceptable gRNAs were available, up to 12 gRNAs were included.

### Next Generation Sequencing and Analysis

The extracted plasmids were amplified using “lib-seq” primers in the Supplementary Table 6 for 10 cycles using Phusion PCR kit (NEB). Each 50 μL PCR reaction consisted of 6 μL of plasmid library, 10 μL of 5X Phusion HF buffer, 0.5 μL of Phusion DNA Polymerase, 10 μM Forward Primer, 10 μM Reverse Primer, and 10 mM dNTPs. Three replicate reactions per sample were amplified. The PCR conditions were: 98°C for 1 minute for initial denaturing, then 98°C for 18s, 66°C for 18s, and 72°C for 30s for 10 cycles, then 72°C for 10 minutes. The products were purified using AMPure XP beads (Beckman-Coulter, Brea, CA, USA). The Illumina adapters and the indices were added by a second PCR using Q5 polymerase (NEB). Each 50 μL PCR consisted of 2-10 μL of purified PCR from the first PCR, 10 μL of 5X Q5 Reaction Buffer, 10 μL 5X Q5 High GC Enhancer, 0.5 μL of Q5 High-Fidelity DNA Polymerase, 10 μM Forward Primer, 10 μM Reverse Primer, 10 mM dNTPs. The PCRs were done in two steps. First step conditions were: 98°C for 1 minute for initial denaturing, then 98°C for 18s, 62°C for 18s, and 72°C for 30s for 4 cycles. Second stage of the PCR was 98°C for 18s, 66°C for 18s, and 72°C for 30s for variable cycles and finally 72°C for 10 minutes. The number of cycles should be determined using a parallel qPCR to ensure that the sample does not saturate. The samples were purified using AMPure XP beads. The samples were run on an Illumina Hi-Seq 4000 for 75 cycles paired-end with a 58-17 breakdown for read 1 and read 2. We used Cutadapt and Bowtie2 to trim the sequences and map them to the targeting sequences with a maximum of one mismatch (43,44). We calculated the log frequency of the reads as:

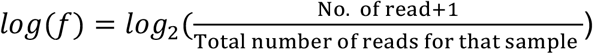

To calculate the depletion scores, we filtered the gRNAs to have a minimum read frequency of 10^−5^ in either the treatment or the control group, e.g. they needed to have a minimum number of reads in the induced or the uninduced sample. The frequency of reads, the number of gRNAs with a minimum of 1 read, and the number of gRNAs passing the threshold are depicted in Supplementary Figure 1. When a gRNA is common to two genes, we assigned its effect to both genes. The distribution of gRNAs per gene in each sample is shown in Supplementary Figure 3. The gRNA depletion/enrichment score between sample A and B was calculated using:

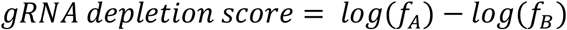

To assign a gene depletion or enrichment score, we sorted the gene log-fold change values of the gRNAs targeting a gene. Then we averaged the highest top three values and also bottom three values. We compared the absolute numbers for the top three and the bottom three gRNAs targeting any given gene and we assigned the largest absolute value as the depletion/enrichment score for that gene. If there were less than six but more than three gRNAs present for a gene, for example g_1-4_ sorted based on their values, we compared the average for g_1-3_ and g_2-4_. For genes with three or less gRNAs, we assigned the average of the gRNAs as the gene depletion score. Previous studies (23) used a metric of average phenotype for the top three most effective gRNAs for each gene. The method presented here has the benefit that it would account for the possibility that repression of some genes could, in fact, be beneficial for growth.

To create the synthetic scrambled genes, we utilized the gRNA depletion scores of the synthetic randomly shuffled gRNAs. We simulated a pool of synthetic scrambled genes by sampling the depletion scores of the synthetic randomly shuffled gRNAs, while maintaining the distribution of gRNA per gene in the sequenced CRISPRi library. Synthetic scrambled genes were populated by considering each yeast ORF and replacing its corresponding gRNAs depletion scores from the pool of synthetic randomly shuffled gRNAs. This process was repeated 200 times to generate a distribution of synthetic scrambled genes. The gene depletion scores were converted to Z scores based on the distribution of synthetic scrambled genes in that replicate. The gene scores (Z-Scores) for the replicates were then averaged using Stouffer’s method.

We used the depletion z Score to classify each gene. For a given z Score threshold, we considered genes with depletion scores below that threshold to be hits (e.g. dosage sensitive). We assessed True Positives based on the genes known to be in a given category (e.g. haploinsufficient or essential genes) and False Positives based on the pool of synthetic scrambled genes. True positive rate (TPR) and false positive rates (FPR) were calculated for a range of z Score thresholds and the Area Under the ROC Curve was assessed. Given the uneven numbers of positives and negatives, we calculated the FDR for each decision value (threshold) using:

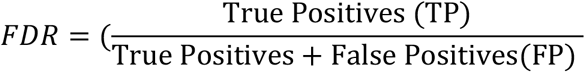

where we oversampled the positive genes.

### IPAGE

We ran iPAGE (29) in continuous mode on the average depletions/enrichment scores of guide RNAs calculated at the gene level as described above. iPAGE discovers gene categories that are significantly informative (*p-*value<0.05) of the average scores. Scores were sorted in 7 bins and only pathway categories were considered.

## Availability of data and materials

All data generated or analyzed during this study are included in the Supplementary Tables associated with this manuscript (See Supplementary Information for details). Accession codes for all sequencing data will be available before publication. The CRISPRi library will be deposited on Addgene. The custom developed code will be provided upon reasonable request.

## Acknowledgements

We thank members of the Tavazoie laboratory for helpful discussions and feedback on the manuscript, specially Wenyan Jiang and Balaji Santhanam. AMR was supported by a Ruth Kirschstein NRSA Postdoctoral Fellowship Award (F32-GM125170) from NIGMS. ST was supported by grants from the NIH (R01-AI077562 and R01-HG009065).

## Contributions

AMR conceived the study and designed the experiments with the help of PO and ST. AMR performed the experiments with the help of PO and MZ. AMR and PO analyzed the results. AMR, PO and ST wrote the manuscript. All authors read and approved the final manuscript.

---

**Supplementary Figure 1.**
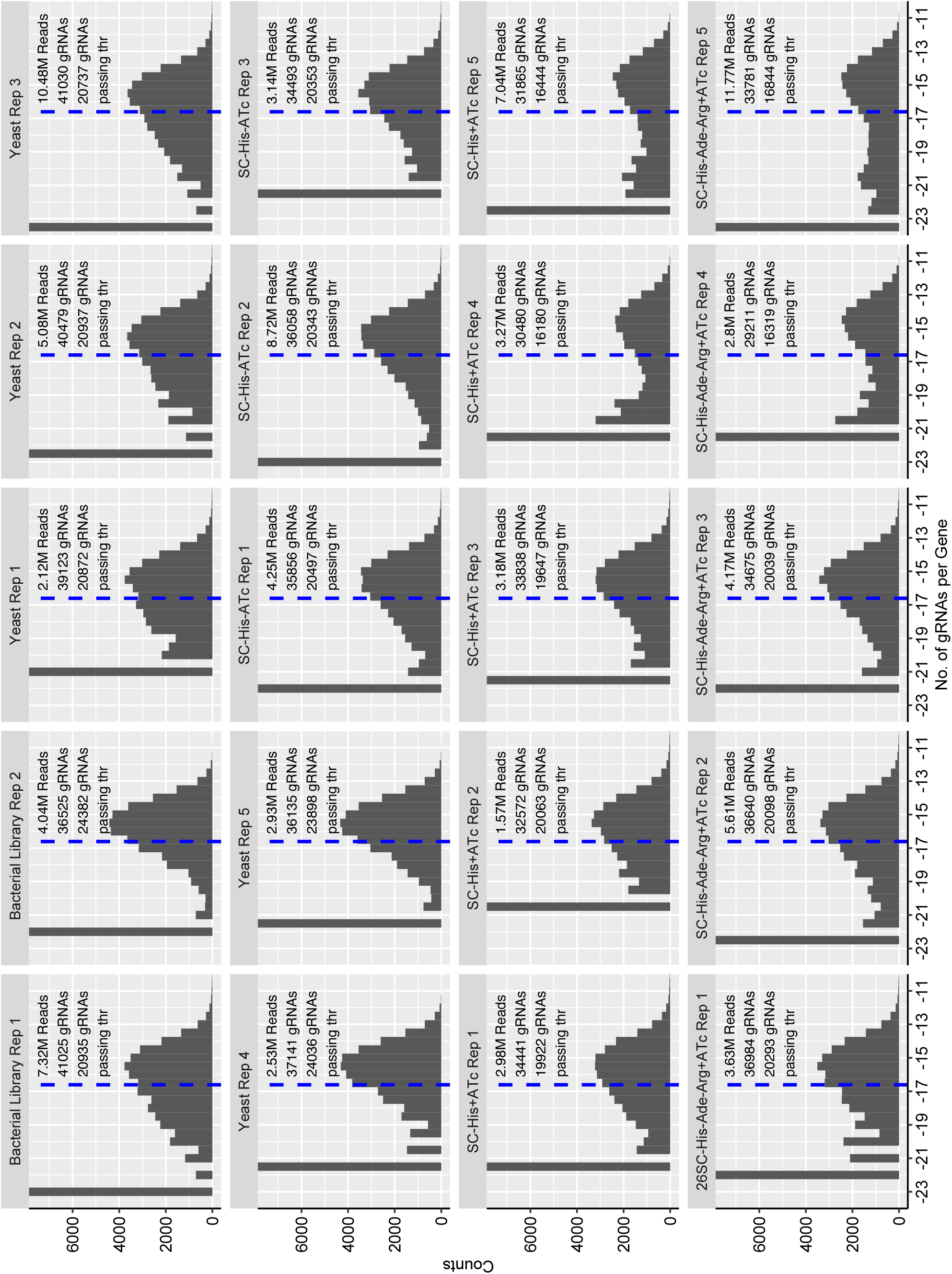
Histograms depicting the frequency of gRNA read distributions in each sample. The total number of reads and gRNAs present for each sample, and the number of gRNAs with read frequency above the threshold is reported accordingly. Dashed line corresponds to the minimum read frequency threshold ~ log2(10^−5^).

**Supplementary Figure 2.**
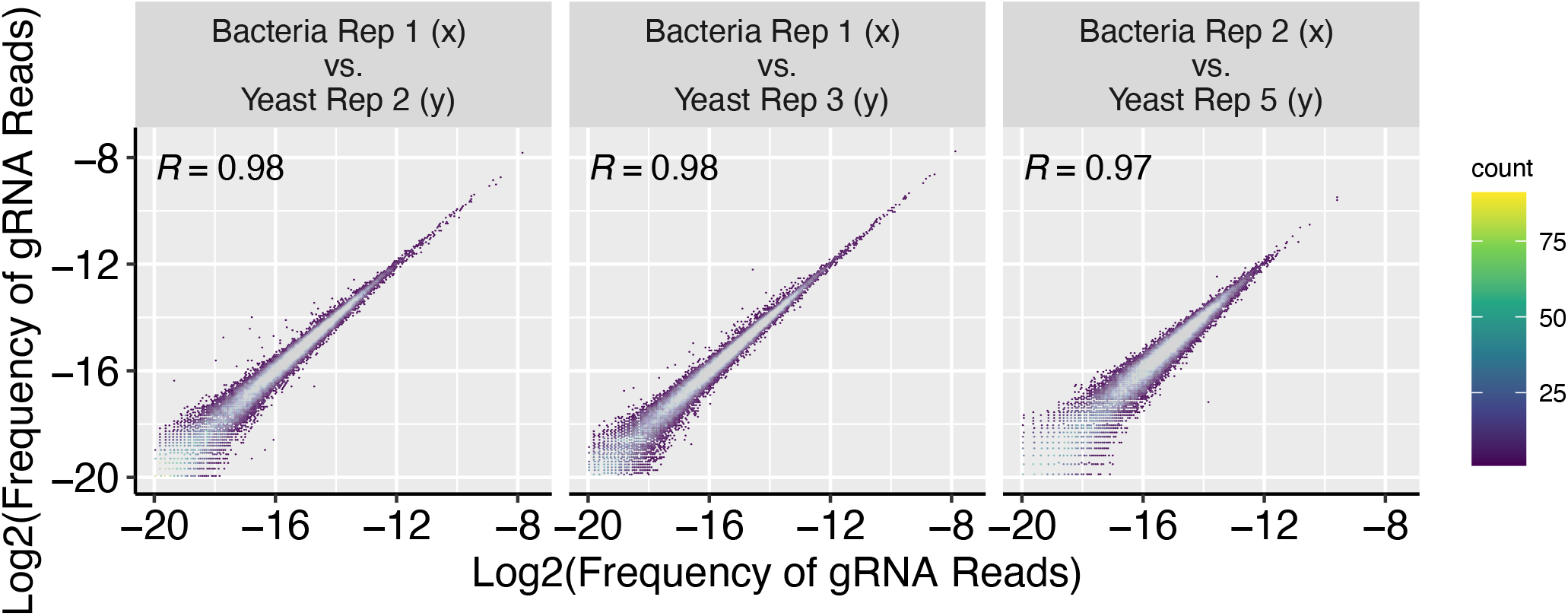
Scatter plot depicting the frequency of reads per gRNA between biological replicates of the CRISPRi library. Pearson Correlation R value is reported for each pair.

**Supplementary Figure 3.**
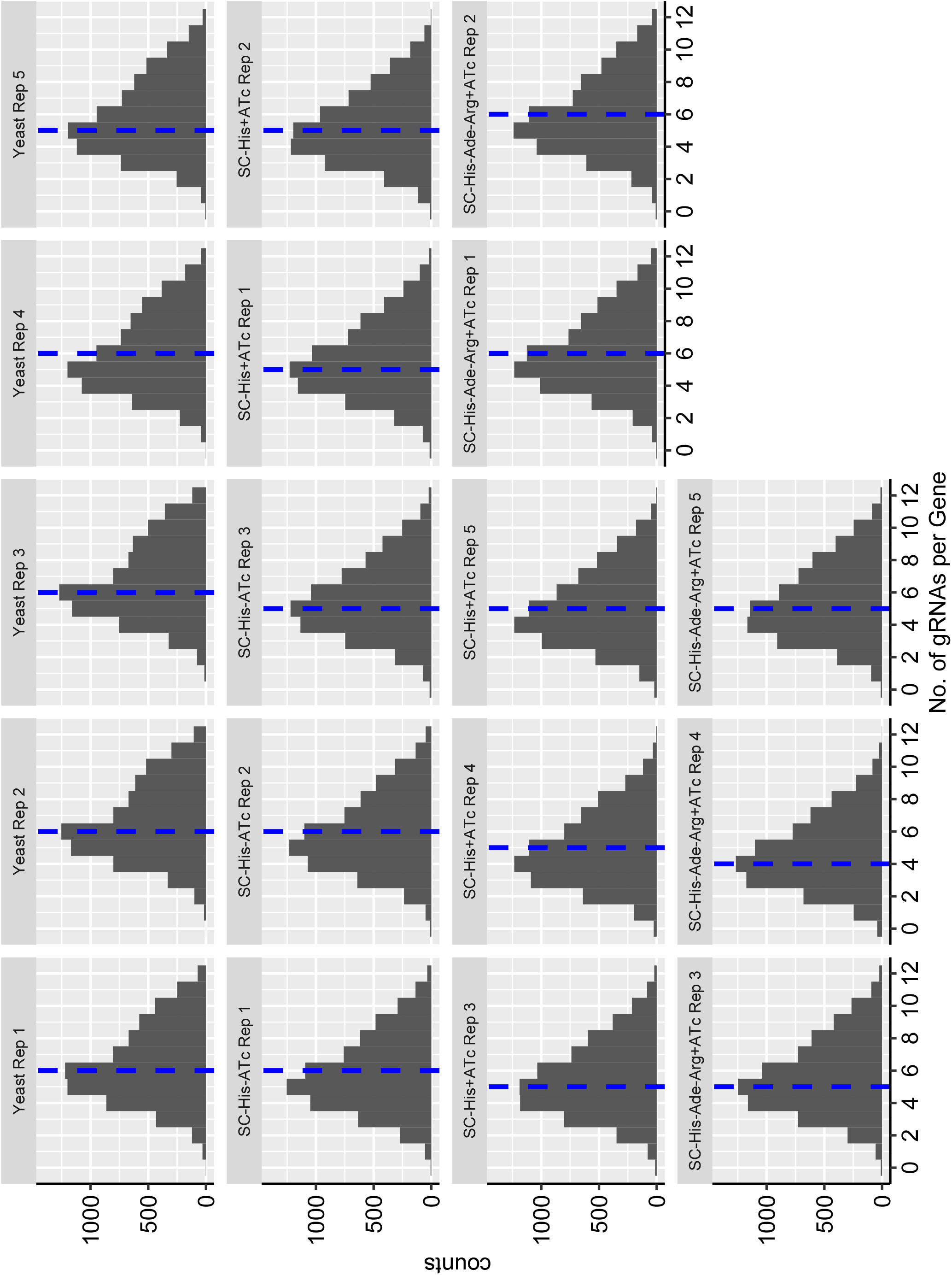
Histogram depicting the number of gRNAs per gene in different samples. The dashed blue line denotes the median. Yeast Rep 1-5 are the yeast transformants.

**Supplementary Figure 4.**
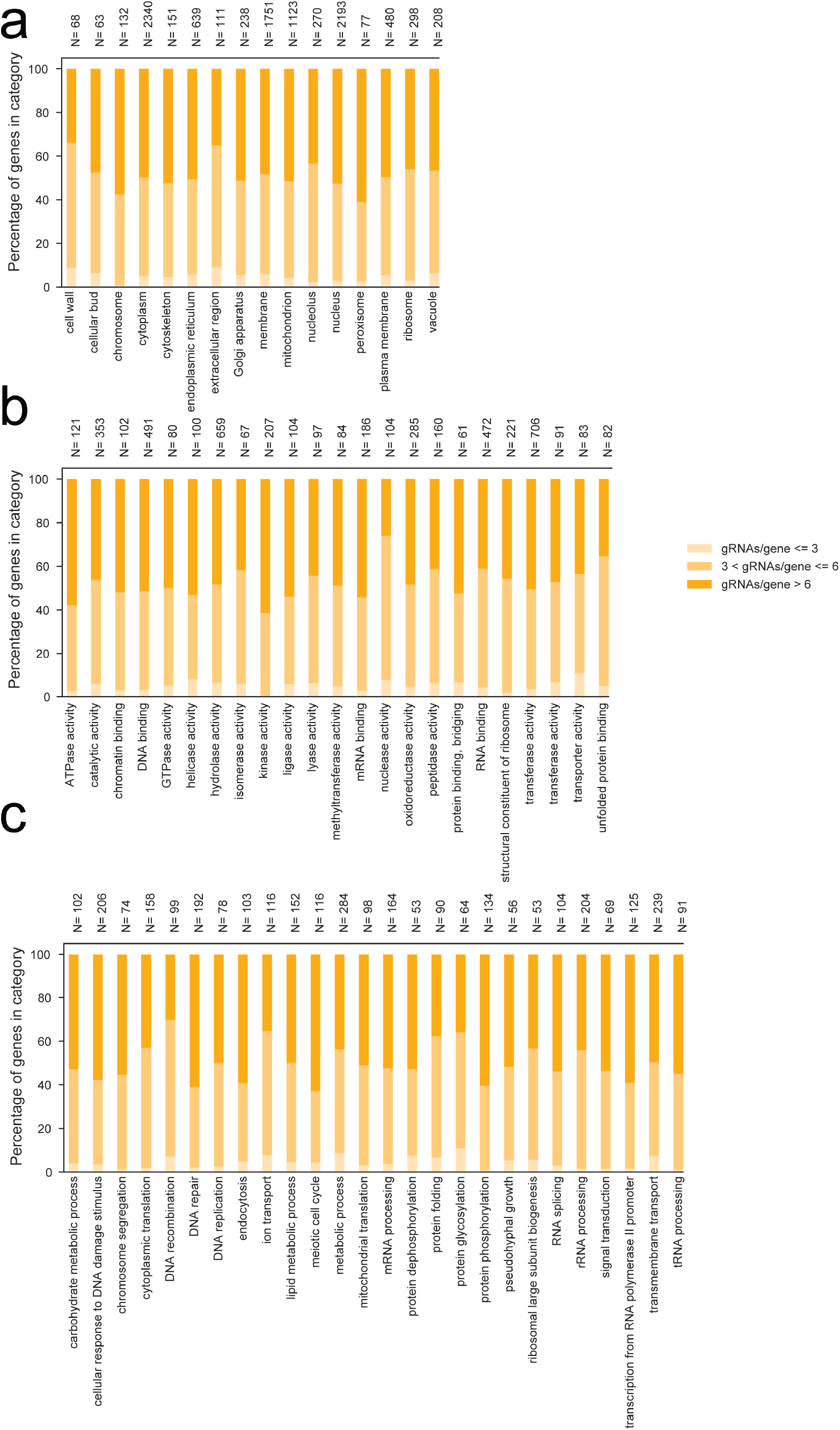
**a)** The distribution of gRNAs per gene across different compartments does not show any bias. **b)** The distribution of gRNAs per gene across different molecular functions shows a good representation. **c)** The distribution of gRNAs per gene across different biological processes does not show any bias.

**Supplementary Figure 5.**
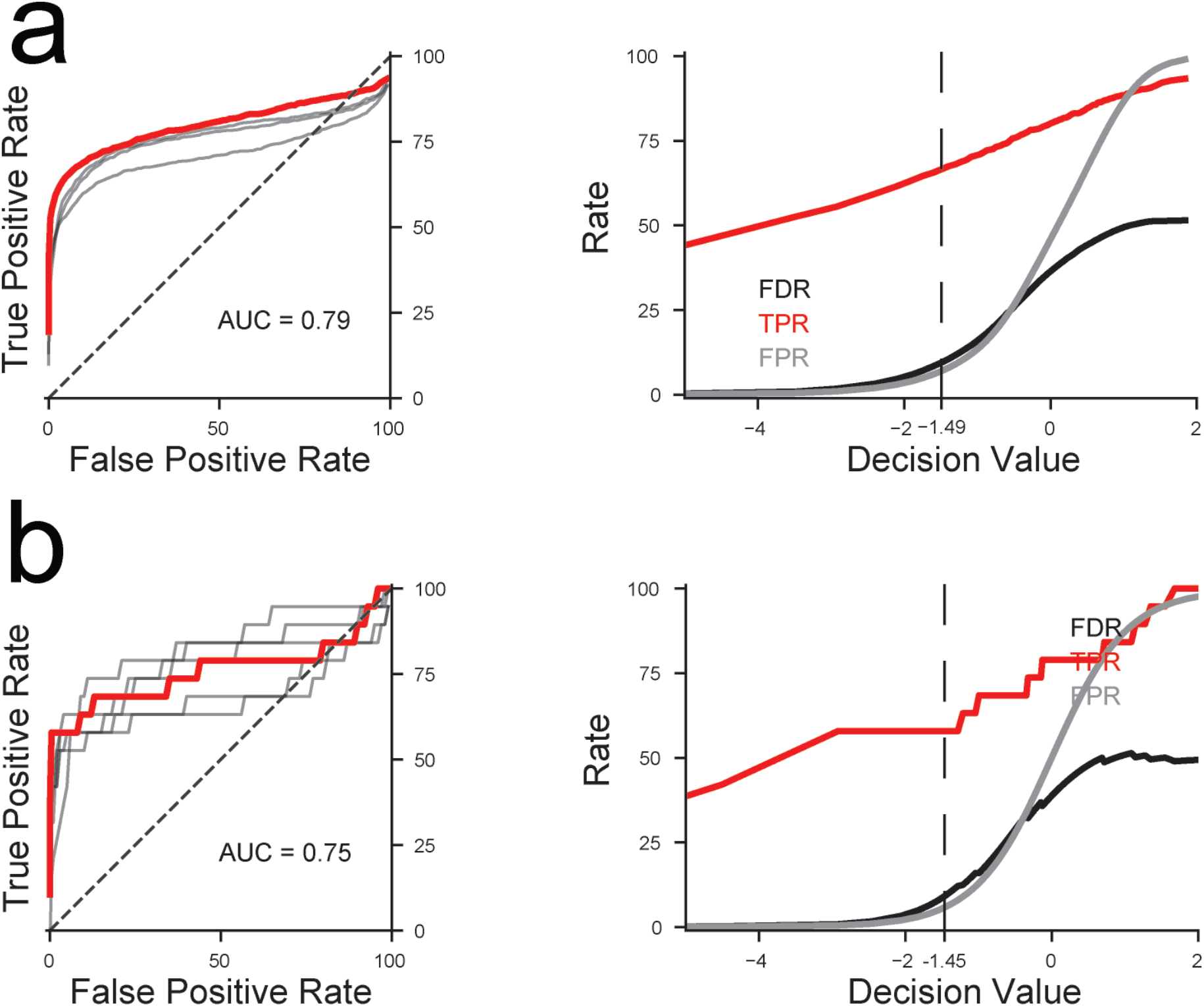
**a)** ROC curve analysis for the detection of essential genes and FDR, TPR and FPR trends based on decision values. ROC curve shows that the depletion Z Score is a good classifier for essential genes. The individual replicates are shown in grey. Area under the curve is 0.79. Dashed line denotes the decision value for FDR<10%. **b)** ROC curve for the detection of adenine and arginine biosynthesis genes and FDR, TPR and FPR trends based on decision values. ROC curve shows that the depletion Z Score is a good classifier for adenine and arginine biosynthetic genes. The individual replicates are shown in grey. Area under the curve is 0.75. Dashed line denotes the decision value for FDR<10%.

**Supplementary Figure 6.**
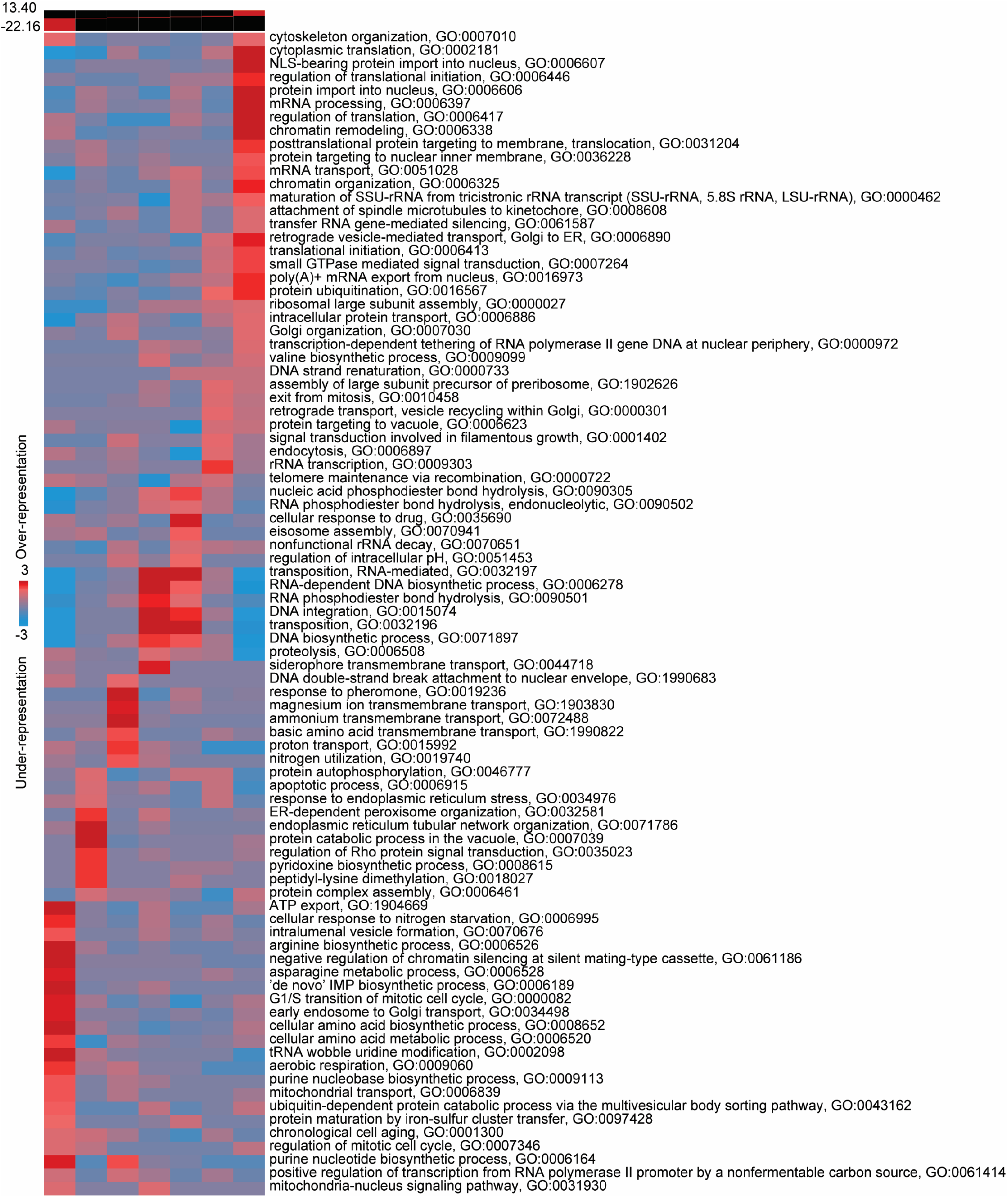
iPAGE pathway analysis results shows all the biological processes that are perturbed in a statistically significant manner as a result of adenine and arginine starvation.

## Supplementary Tables

**Table 1**. Summary of the designed spacers.

**Table 2**. Summary of the control synthetic randomly shuffled spacers.

**Table 3**. Summary of frequency of reads in each sample.

**Table 4. Sheet (1)** Gene depletion scores for samples with ATc versus without ATc. **Sheet (2)** Gene depletion scores for samples in the drop-out media for Histidine, Arginine and Adenine versus Histidine drop-out media. **Sheet (3)** Gene depletion scores for samples in the drop-out media for Histidine, Arginine and Adenine versus Histidine drop-out media for gRNAs with PAM between 0 and 150 bp upstream of TSS.

**Table 5**. List of the gene categories used in this study

**Table 6**. List of the primer sequences used in this study

## References

1. Burns N, Grimwade B, Ross-Macdonald PB, Choi EY, Finberg K, Roeder GS, et al. Large-scale analysis of gene expression, protein localization, and gene disruption in Saccharomyces cerevisiae. Genes & Development. Cold Spring Harbor Lab; 1994 May 1;8(9):1087–105.

2. Deutschbauer AM, Jaramillo DF, Proctor M, Kumm J, Hillenmeyer ME, Davis RW, et al. Mechanisms of haploinsufficiency revealed by genome-wide profiling in yeast. Genetics. 2005 Apr;169(4):1915–25.

3. Giaever G, Shoemaker DD, Jones TW, Liang H, Winzeler EA, Astromoff A, et al. Genomic profiling of drug sensitivities via induced haploinsufficiency. Nat Genet. 1999 Mar;21(3):278–83.

4. Girgis HS, Liu Y, Ryu WS, Tavazoie S. A comprehensive genetic characterization of bacterial motility. PLoS Genet. 2007 Sep;3(9):1644–60.

5. Qian W, Ma D, Xiao C, Wang Z, Zhang J. The genomic landscape and evolutionary resolution of antagonistic pleiotropy in yeast. Cell Rep. 2012 Nov 29;2(5):1399–410.

6. Smith V, Botstein D, Brown PO. Genetic footprinting: a genomic strategy for determining a gene's function given its sequence. Proc Natl Acad Sci USA. National Academy of Sciences; 1995 Jul 3;92(14):6479–83.

7. Winzeler EA, Shoemaker DD, Astromoff A, Liang H, Anderson K, Andre B, et al. Functional characterization of the S. cerevisiae genome by gene deletion and parallel analysis. Science. 1999 Aug 6;285(5429):901–6.

8. Jinek M, Chylinski K, Fonfara I, Hauer M, Doudna JA, Charpentier E. A programmable dual-RNA-guided DNA endonuclease in adaptive bacterial immunity. Science. American Association for the Advancement of Science; 2012 Aug 17;337(6096):816–21.

9. Cong L, Ran FA, Cox D, Lin S, Barretto R, Habib N, et al. Multiplex genome engineering using CRISPR/Cas systems. Science. 2013 Feb 15;339(6121):819–23.

10. DiCarlo JE, Norville JE, Mali P, Rios X, Aach J, Church GM. Genome engineering in Saccharomyces cerevisiae using CRISPR-Cas systems. Nucleic Acids Research. 2013 Apr;41(7):4336–43.

11. Sidik SM, Huet D, Ganesan SM, Huynh M-H, Wang T, Nasamu AS, et al. A Genome-wide CRISPR Screen in Toxoplasma Identifies Essential Apicomplexan Genes. Cell. 2016 Sep 8;166(6):1423–1435.e12.

12. Shalem O, Sanjana NE, Hartenian E, Shi X, Scott DA, Mikkelson T, et al. Genome-scale CRISPR-Cas9 knockout screening in human cells. Science. 2014 Jan 3;343(6166):84–7.

13. Wang T, Wei JJ, Sabatini DM, Lander ES. Genetic screens in human cells using the CRISPR-Cas9 system. Science. 2014 Jan 3;343(6166):80–4.

14. Bassett AR, Kong L, Liu J-L. A genome-wide CRISPR library for high-throughput genetic screening in Drosophila cells. J Genet Genomics. 2015 Jun 20;42(6):301–9.

15. Bikard D, Jiang W, Samai P, Hochschild A, Zhang F, Marraffini LA. Programmable repression and activation of bacterial gene expression using an engineered CRISPR-Cas system. Nucleic Acids Research. 2013 Aug;41(15):7429–37.

16. Qi LS, Larson MH, Gilbert LA, Doudna JA, Weissman JS, Arkin AP, et al. Repurposing CRISPR as an RNA-guided platform for sequence-specific control of gene expression. Cell. 2013 Feb 28;152(5):1173–83.

17. Gilbert LA, Larson MH, Morsut L, Liu Z, Brar GA, Torres SE, et al. CRISPR-Mediated Modular RNA-Guided Regulation of Transcription in Eukaryotes. Cell. Elsevier Inc; 2013 Jul 18;154(2):442–51.

18. Farzadfard F, Perli SD, Lu TK. Tunable and multifunctional eukaryotic transcription factors based on CRISPR/Cas. ACS Synth Biol. 2013 Oct 18;2(10):604–13.

19. Perez-Pinera P, Kocak DD, Vockley CM, Adler AF, Kabadi AM, Polstein LR, et al. RNA-guided gene activation by CRISPR-Cas9-based transcription factors. Nat Meth. 2013 Oct;10(10):973–6.

20. Dominguez AA, Lim WA, Qi LS. Beyond editing: repurposing CRISPR–Cas9 for precision genome regulation and interrogation. Nature Reviews Molecular Cell Biology. Nature Publishing Group; 2015 Dec 16;17(1):5–15.

21. Chavez A, Scheiman J, Vora S, Pruitt BW, Tuttle M, P R Iyer E, et al. Highly efficient Cas9-mediated transcriptional programming. Nat Meth. 2015 Mar 2;12(4):326–8.

22. Smith JD, Suresh S, Schlecht U, Wu M, Wagih O, Peltz G, et al. Quantitative CRISPR interference screens in yeast identify chemical-genetic interactions and new rules for guide RNA design. Genome Biol. Genome Biology; 2016 Mar 7;:1–16.

23. Gilbert LA, Horlbeck MA, Adamson B, Villalta JE, Chen Y, Whitehead EH, et al. Genome-Scale CRISPR-Mediated Control of Gene Repression and Activation. Cell. Elsevier Inc; 2014 Oct 23;159(3):647–61.

24. Smith JD, Schlecht U, Xu W, Suresh S, Horecka J, Proctor MJ, et al. A method for high-throughput production of sequence-verified DNAlibraries and strain collections. Molecular Systems Biology. John Wiley & Sons, Ltd; 2017 Feb 9;13(2):913–5.

25. Lian J, Schultz C, Cao M, HamediRad M, Zhao H. Multi-functional genome-wide CRISPR system for high throughput genotype-phenotype mapping. Nature Communications. Nature Publishing Group; 2019 Dec 19;10(1):5794–10.

26. Hoon S, Smith AM, Wallace IM, Suresh S, Miranda M, Fung E, et al. An integrated platform of genomic assays reveals small-molecule bioactivities. Nat Chem Biol. 2008 Aug;4(8):498–506.

27. Gibson DG, Young L, Chuang R-Y, Venter JC, Hutchison CA, Smith HO. Enzymatic assembly of DNA molecules up to several hundred kilobases. Nat Meth. 2009 May;6(5):343–5.

28. Gietz RD, Schiestl RH. High-efficiency yeast transformation using the LiAc/SS carrier DNA/PEG method. Nat Protoc. 2007;2(1):31–4.

29. Goodarzi H, Elemento O, Tavazoie S. Revealing Global Regulatory Perturbations across Human Cancers. Molecular Cell. Elsevier Ltd; 2009 Dec 11;36(5):900–11.

30. Crabeel M, Soetens O, De Rijcke M, Pratiwi R, Pankiewicz R. The ARG11 gene of Saccharomyces cerevisiae encodes a mitochondrial integral membrane protein required for arginine biosynthesis. J Biol Chem. American Society for Biochemistry and Molecular Biology; 1996 Oct 4;271(40):25011–8.

31. Jauniaux JC, Urrestarazu LA, bacteriology JWJO, 1978. Arginine metabolism in Saccharomyces cerevisiae: subcellular localization of the enzymes. Am Soc Microbiol

32. Jones E, Fink G. Regulation of amino acid and nucleotide biosynthesis in yeast. The Molecular Biology of the Yeast Saccharomyces Metabolism And Gene Expression. 1982;:181–299.

33. Ljungdahl PO, Daignan-Fornier B. Regulation of amino acid, nucleotide, and phosphate metabolism in Saccharomyces cerevisiae. Genetics. Genetics; 2012 Mar;190(3):885–929.

34. Messenguy F. Regulation of arginine biosynthesis in Saccharomyces cerevisiae: isolation of a cis-dominant, constitutive mutant for ornithine carbamoyltransferase synthesis. Journal of Bacteriology. American Society for Microbiology Journals; 1976 Oct 1;128(1):49–55.

35. Myasnikov AN, Sasnauskas KV, Janulaitis AA, Smirnov MN. The Saccharomyces cerevisiae ADE1 gene: structure, overexpression and possible regulation by general amino acid control. Gene. 1991 Dec 20;109(1):143–7.

36. Tibbetts AS, Appling DR. Saccharomyces cerevisiae expresses two genes encoding isozymes of 5-aminoimidazole-4-carboxamide ribonucleotide transformylase. Arch Biochem Biophys. 1997 Apr 15;340(2):195–200.

37. Michaillat L, Mayer A. Identification of genes affecting vacuole membrane fragmentation in Saccharomyces cerevisiae. PLoS ONE. 2013;8(2):e54160.

38. Caldovic L, Tuchman M. N-acetylglutamate and its changing role through evolution. Biochem J. 2003 Jun 1;372(Pt 2):279–90.

39. Messenguy F, Dubois E. Role of MADS box proteins and their cofactors in combinatorial control of gene expression and cell development. Gene. 2003 Oct 16;316:1–21.

40. Forsberg H, Gilstring CF, Zargari A, Martínez P, Ljungdahl PO. The role of the yeast plasma membrane SPS nutrient sensor in the metabolic response to extracellular amino acids. Mol Microbiol. John Wiley & Sons, Ltd; 2001 Oct;42(1):215–28.

41. Jiang W, Oikonomou P, Tavazoie S. Comprehensive Genome-wide Perturbations via CRISPR Adaptation Reveal Complex Genetics of Antibiotic Sensitivity. Cell. 2020 Feb 21.

42. Gueldener U, Heinisch J, Koehler GJ, Voss D, Hegemann JH. A second set of loxP marker cassettes for Cre-mediated multiple gene knockouts in budding yeast. Nucleic Acids Research. 2002 Mar 15;30(6):e23–3.

43. Martin M. Cutadapt removes adapter sequences from high-throughput sequencing reads. EMBnetjournal. 2011 May 2;17(1):10–2.

44. Langmead B, Salzberg SL. Fast gapped-read alignment with Bowtie 2. Nat Meth. 2012 Mar 4;9(4):357–9.

